# The design, analysis and application of mouse clinical trials in oncology drug development

**DOI:** 10.1101/425256

**Authors:** Sheng Guo, Xiaoqian Jiang, Binchen Mao, Qi-Xiang Li

**Affiliations:** Crown Bioscience Inc., 218 Xinghu Street, Suzhou Industrial Park, Jiangsu, China, 215028.; Crown Bioscience, Inc., 3375 Scott Blvd, Suite 108, Santa Clara, CA 95054, USA.; State Key Laboratory of Natural and Biomimetic Drugs, Peking University, Beijing, China, 100191.

**Keywords:** PDX, CDX, syngeneic model, mouse clinical trials, linear mixed models, survival analysis, statistical power, biomarker

## Abstract

Mouse clinical trials (MCTs) are becoming widely used in pre-clinical oncology drug development. In this study, we provide some general guidelines on the design, analysis and application of MCTs. We first established empirical quantitative relationships between mouse number and measurement accuracy for both categorical and continuous efficacy endpoints, and showed that more mice are needed to achieve given accuracy for syngeneic models than for PDXs and CDXs. There is considerable disagreement between categorical methods on calling drug responses as objective response, indicating limitations of such approaches. We then introduced linear mixed models, or LMMs, to describe MCTs as clustered longitudinal studies, which explicitly model growth and drug response heterogeneities across mouse models and among mice within a mouse model. Several case studies were used to demonstrate the advantages of LMMs in discovering biomarkers and exploring a drug’s mechanism of action. We also introduced the additive frailty models to perform survival analysis on MCTs, which more accurately estimate hazard ratios by modeling the clustered population structures in MCTs. We performed computational simulations for LMMs and frailty models to generate statistical power curves, and showed that statistical power is close for designs with similar total number of mice at given drug efficacy. Finally, we explained how MCTs can explain discrepant results in clinical trials, hence, MCTs are more than preclinical versions of clinical trials but possess their unique values. Results in the report will make MCTs a better tool for oncology drug development.

## Introduction

Cancer is a heterogeneous disease with intra- and inter-tumor genomic diversity that determines cancer initiation, progression and treatment. The understandings of cancer biology and the development of therapeutics have been aided greatly by a variety of mouse tumor models, including cell line-derived xenografts (CDXs), patient derived-xenografts (PDXs), genetically engineered mouse models (GEMMs), cell line- or primary tumor-derived homografts in syngeneic mice and so on (reviewed by (Day, Merlino, & Van Dyke, 2015; Khaled & Liu, 2014; Li, Feuer, Ouyang, & An, 2017; Walrath, Hawes, Van Dyke, & Reilly, 2010)). These models differ in their generation, host and tumor genomics and biology, availability, and research utilizations. For example, immunotherapies are tested in immunocompetent models such as GEMMs and syngeneic models.

Past decades witnessed the accelerated creation, distribution, profiling and characterization of mouse tumor models (Byrne et al., 2017; Gao et al., 2015; Guo et al., 2016; Krupke et al., 2017; Stewart et al., 2017; Townsend et al., 2016). The abundant collections made it possible to conduct the so-called “mouse clinical trials (MCTs)”, in which a panel of mouse models, dozens to hundreds, are used to evaluate therapeutic efficacy, discover/validate biomarkers, study tumor biology and so on. MCTs demonstrated faithful clinical predictions in multiple studies (Bardelli et al., 2013; Bertotti et al., 2011; Bertotti et al., 2015; Gao et al., 2015; Migliardi et al., 2012; Yao et al., 2017). While most reported MCTs used PDXs, MCTs using other mouse models, such as syngeneic models, are now widely performed as well.

Because of their resemblance to clinical trials, MCTs are often analyzed by methods for clinical trials. For example, overall survival (OS) and progression-free survival (PFS) are estimated by tumor volume increase, Cox proportional hazards models are used for survival analysis, response categories are defined by tumor volume change and objective response rate (ORR) is calculated (Bertotti et al., 2015; Gao et al., 2015; Houghton et al., 2007). However, MCTs differ from clinical trials in many ways: (1) a mouse model have mice in all arms, one is usually a vehicle arm; (2) there can be more than one mouse for each mouse model in each arm; (3) tumor volumes are routinely measured; (4) mouse models are usually characterized with genomic/pharmacology/histopathology annotations; (5) MCTs are done in labs that reduces/removes various noise and inconvenience encountered in clinical trials, such as dropouts, long trial time and concomitant medication.

In this study, we combine empirical data analysis, statistical modeling and computational simulations to address some key issues for MCTs, including the determination of animal numbers (number of mouse models and number of mice per mouse model), statistical power calculation, quantification of efficacy difference between mice/mouse models/drugs, survival analysis, biomarker discovery/validation with and beyond simple efficacy readouts, handling of mouse dropouts, missing data and difference in tumor growth rates, study of mechanisms of action (MoA) for drugs. We will also show MCTs can explain discrepant clinical trial results.

## RESULTS

### Determining number of mice for categorical responses

We collected tumor volume data under drug treatment for 26127 mice from 2883 unique treatment PDXs, 11139 mice from 1219 unique treatment CDXs, and 5945 mice from 637 unique treatment syngeneic models. A unique treatment model is a mouse model treated by a drug in a study. Every unique treatment has at least 8 mice. Categorical drug response was determined by 4 methods (see Materials and Methods), and we illustrate the results using the mRECIST criteria, which classifies drug response into 4 categories: complete response (CR), partial response (PR), stable disease (SD), and progressive disease (PD). For each unique treatment model, its response is the majority response of all mice. We observed that individual mouse responses matched the majority response most often for PD: 90% for PDXs, 95% for CDXs and syngeneic models (Fig. 1a-c). The other 3 response categories exhibit lower concordance, particularly so for syngeneic models. Of the 10 unique treatment syngeneic models classified as CR, only half of the mice had complete response as well, while 17% of mice were PD and resistant to treatment. Such polarized response pattern is observed in the other 3 methods, too (**Fig. S1-S3**). Large variance exists for all 4 response categories. For example, only about 70% of individual responses matched the majority response for a third of the 107 unique treatment PDX models categorized as CR, although the average is 83%.

**Figure 1.**
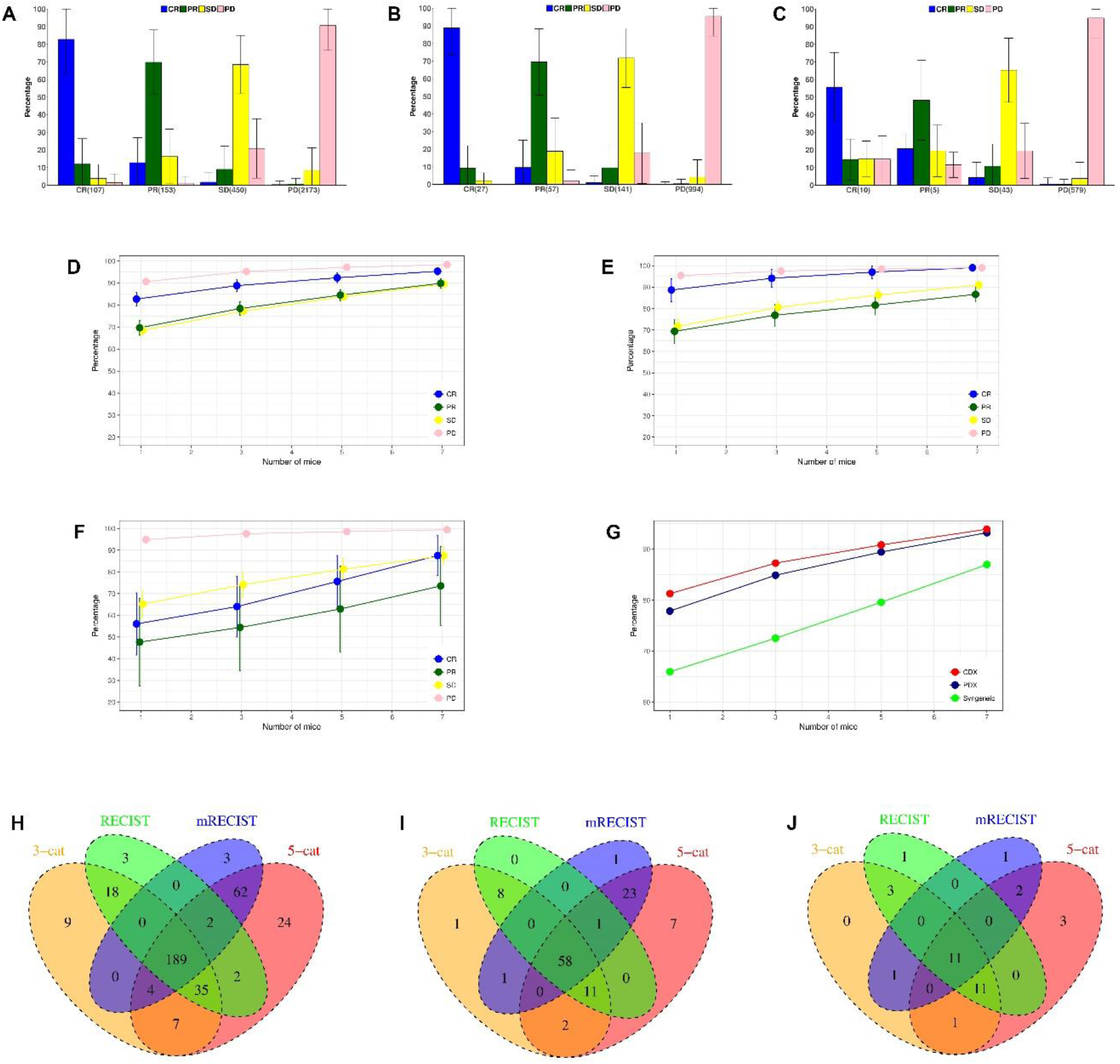
Mouse number and measurement accuracy of categorical responses defined by the mRECIST criteria. (**a-c**): individual mouse response and majority response in PDX (**a**), CDX (**b**) and syngeneic models (**c**), x axis is the number of majority response from 4 response categories (CR: complete response, PR: partial response, SD: stable disease, PD: progressive disease.), y axis is the percentage of individual mouse response relative to the majority (average ± s.d.). There are 26,127 mice in 2,883 unique treatment PDX models, 11,139 mice in 1,219 unique treatment CDX models, and 5,945 mice in 637 unique treatment syngeneic models. Each unique treatment model had at least 8 mice. (**d-g**): measurement accuracy increases with number of mice for PDX (**d**), CDX (**e**) and syngeneic models (**f**). For each unique treatment model, the majority response of n (n=1, 3, 5, 7 in x axis) randomly sampled mice was obtained to see if it agreed with the actual majority response. The procedure was repeated 1,000 times to obtain the accuracy—percentage of times (average ± s.d.) that they agreed—for the 4 response categories, whose unweighted average is shown in (**g**). (h-j): Venn diagram showing the overlap of unique treatment PDX models classified as objective response by 4 categorical methods in PDX (**h**), CDX (**i**), and syngeneic models (**j**). Objective response is OR in the 3-cat method, CR+PR in the mRECIST and RECIST methods, MCR+CR+PR in the 5-cat method.

Measurement accuracy increases with number of mice. We randomly sampled n (n=1, 3, 5, 7) mice from all the mice in a treatment and obtained a majority response, which was then compared with the actual majority response. The procedure was repeated for 1,000 times to generate statistical results (Fig. 1d-f). Accuracy increases with mouse number for all 4 categories, and their unweighted average is highest in CDXs, which is slightly higher than PDXs, while syngeneic models have much lower accuracy (Fig. 1g). Therefore, more mice are needed for syngeneic models to achieve similar accuracy as PDXs/CDXs. For example, accuracy is comparable between syngeneic studies with 5 mice per model and PDX/CDX studies with 1 mouse per model. Similar patterns are also seen in the other 3 methods (**Fig. S1-S3**).

All the 4 methods categorize responses based on relative tumor volume (RTV) at a later day to treatment initiation day, but differ in specific thresholds. As such, a unique treatment model can be categorized differently. We found that there is a good overlapping for unique treatment models classified as objective response between the 4 methods (Fig. 1h-j), and their objective response rates (ORR) are similar. (**Table S1**). Nevertheless, there are many models only unique to some methods as OR, cautioning method-specific bias and applicability. For example, the mRECIST considers averaging tumor reduction for a period of time, therefore, a unique treatment model can be classified as PD even though tumor completely disappears at end of study (**Fig. S4**).

### Determining number of mice for continuous responses

Drug efficacy can be measured by continuous responses, some are direct adaption of clinical endpoints (e.g., PFS and OS), others are unique to mouse studies that use data from both vehicle and drug treatment groups (e.g., RTV ratio between drug and vehicle groups). We calculated the estimation errors of PFS and RTV ratio computed from n (n=1 to 9) mice randomly sampled from the ≥10 mice in a study, and obtained the quantitative relationship between estimation errors and mouse numbers (Fig. 2). Large estimation errors are inherent to small sample sizes, particularly so for syngeneic models. For example, percent error of PFS is greater than 20% for 63% syngeneic mice and for about half of PDX/CDX mice (Fig. 2d). Estimation errors are reduced sharply by addition of more mice when n is small. For RTV ratio, 3 mice in both drug and vehicle group already lift mice with absolute error <0.2 from 60% to above 80% for PDXs/CDXs (Fig. 2h). Similar results hold for other continuous endpoints as well (**Fig. S5**).

**Figure 2.**
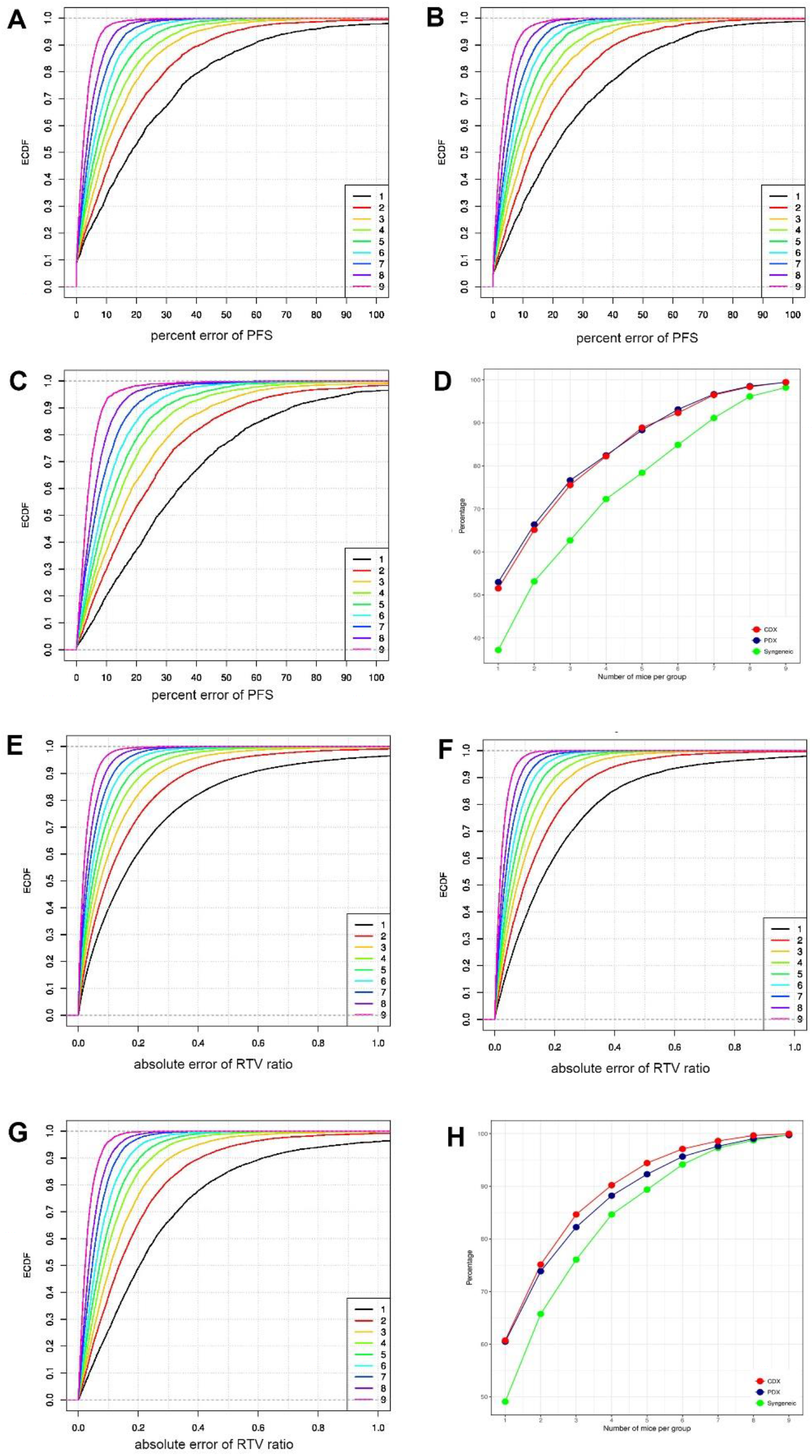
Determining mouse numbers for continuous responses. (**a-c**): Progression-free survival, or PFS, calculated from n mice (n=1 to 9) randomly sampled from a unique treatment m
odel with at least 10 mice shows relative deviation to the PFS calculated from all mice in PDX (**a**), CDX (**b**), and syngeneic models (**c**), x axis is the percent error of PFS, and y axis is the empirical cumulative density function (ECDF) estimated from the random samplings for each n. Percent error of PFS decreases with increased number of mice, and the error is larger for syngeneic models than PDXs/CDXs. (**d**): Percentages of unique treatment models with percent error less than 20% in the 3 types of mouse models. (**e-g**): RTV ratio between drug and vehicle groups, calculated from n mice (n=1 to 9) randomly sample from a study with at least 10 mice in both drug and vehicle groups, shows deviation to the RTV ratio calculated from all mice in both groups in PDX (**e**), CDX (**f**), and syngeneic models (**g**), x axis is the absolute error, and y axis is the empirical cumulative density function (ECDF) estimated from the random samplings for each n. Absolute error of RTV ratio decreases with increased number of mice, and the error is larger for syngeneic models than PDXs/CDXs. (**h**): Percentages of studies with absolute error less than 0.2 in the 3 types of mouse models.

### Modeling MCTs as clustered longitudinal studies

It is convenient to measure drug efficacy by a categorical or continuous endpoint, but those approaches also suffer from loss of information and other drawbacks. For example, it is somewhat arbitrary to choose a day to calculate RTV ratio and TGI; it adds logistic burden to match mice with comparable tumor volume at treatment initiation day (Laajala et al., 2016); it is difficult to deal with mouse dropouts. These shortcomings can be overcome by modeling MCTs as clustered longitudinal studies, in which a cluster is consisted of all mice of a mouse model so they share genomic profile and have more similar drug response. Each mouse is in a longitudinal study. It can be shown that tumor growth in majority of mice follows exponential kinetics (**Fig. S6**). Therefore, we can model the clustered longitudinal studies by a 3-level linear mixed model (LMM) on the log-transformed tumor volumes (logTV) and day (Fig. 3a). There are covariates associated with mouse models such as cancer type and genomic features, which can be used for examining efficacy difference on cancers and for discovering predictive biomarkers.

**Figure 3.**
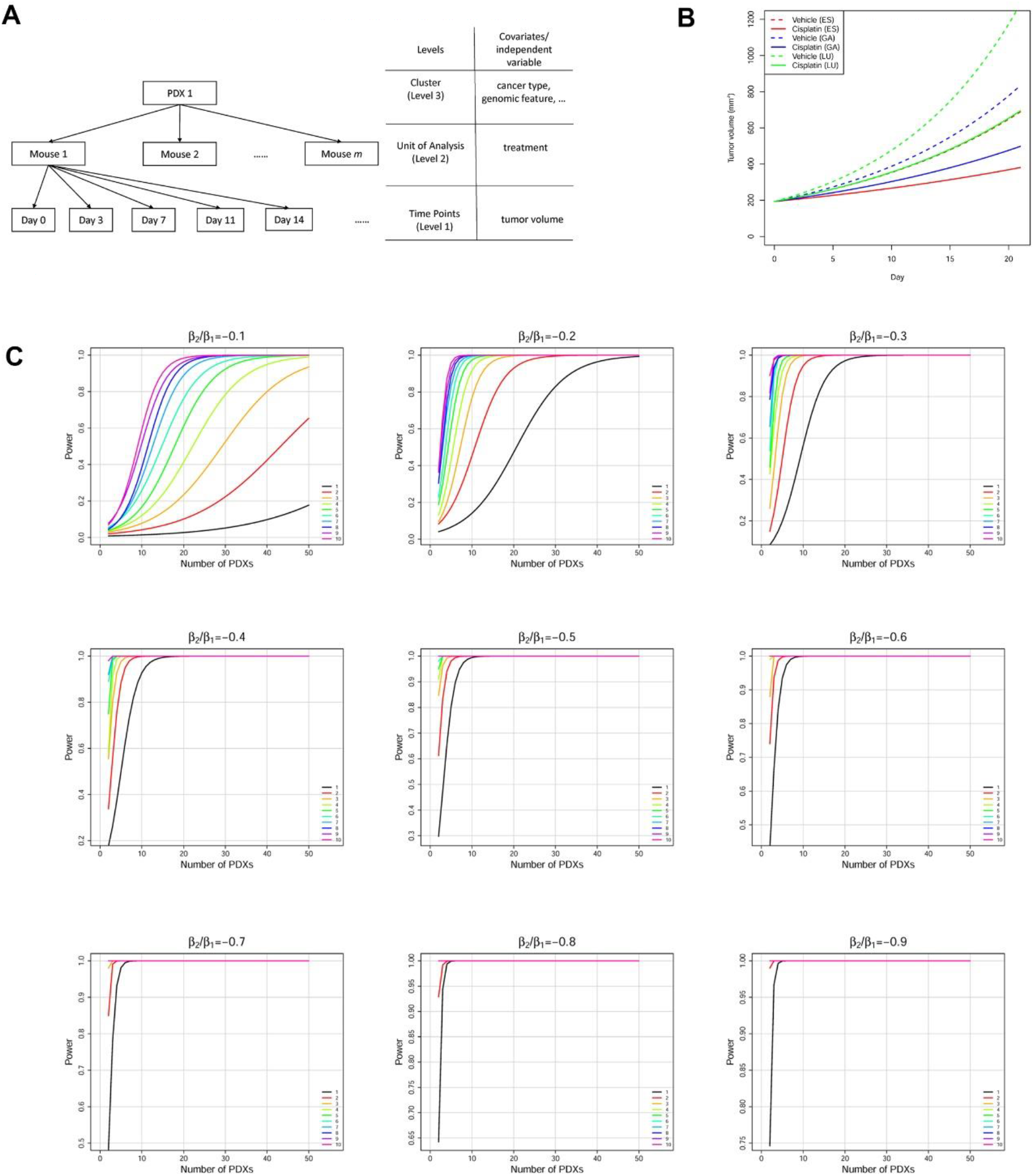
Linear mixed models (LMMs) can be used to model the clustered longitudinal data from MCTs. (**a**) the structure of the clustered longitudinal data for a PDX in a MCT. PDX level and mouse level covariates can be incorporated into LMMs. (**b**) Mean tumor growth curves for 3 cancers under vehicle treatment and cisplatin treatment. (**c**) Statistical power curves of the cisplatin MCT. Power is calculated at significance level α=0.05 when the cisplatin treatment reduces tumor growth rate by 10% to 90%, i.e. β_1_/β_2_= −0.1 to −0.9 in Equation 4 in Materials and Methods. The 10 colored curves in each graph denote the number of mice for every PDX in each arm.

We use one example to demonstrate the modeling of MCTs by LMMs for efficacy evaluation and comparison. In this MCT, cisplatin—a chemotherapy drug—was administrated to 42 PDXs (4mg/kg, weekly dosing for 3 weeks), including 13 esophageal cancers (ES), 21 gastric cancers (GA) and 8 lung cancers (LU), each PDX with 5 to 9 mice (**Fig. S7**). We fit the efficacy data by a LMM (Equation 3 in Materials and Methods), which explicitly models tumor growth rate heterogeneity and drug response heterogeneity at both PDX level and mouse level. Model fitting is satisfactorily (**Fig. S8**, Table 1). We conclude that (1) under vehicle treatment, tumor in GA grows slightly faster than ES, while tumor growth is much faster in LU; (2) cisplatin has comparable efficacy on the 3 cancers (p-values for *β*_5_ and *β*_6_ are >0.05). The results can be readily visualized from the mean growth curves for the 3 cancers under (Fig. 3b).

**Table 1.** Parameters estimated for the LMM (Equation 3) of the cisplatin dataset.

| Fixed-Effect Parameters | Estimate* | p-value |
| --- | --- | --- |
| $\beta_0$ (Intercept) | 5.2641(0.0257) | 0 |
| $\beta_1$ (Day) | 0.0605(0.0043) | 1.5E-43 |
| $\beta_2$ (Day × CancerTypeGA) | 0.0091(0.0055) | 0.098 |
| $\beta_3$ (Day × CancerTypeLU) | 0.0297(0.0071) | 2.8E-5 |
| $\beta_4$ (Day × Treatment) | -0.0282(0.0031) | 1.2E-19 |
| $\beta_5$ (Day × CancerTypeGA × Treatment) | 0.0037(0.0039) | 0.35 |
| $\beta_6$ (Day × CancerTypeLU × Treatment) | -0.0011(0.0052) | 0.84 |

\*parameters estimated by the REML method in the R nlme package.

### Statistical power and sample size determination in MCTs

Much like clinical trials, rational design of MCTs requires statistical power calculation and sample size determination—number of mouse models and number of mice per mouse model. We demonstrate this under the LMM framework with the following assumptions (1) a balanced n:n design in which there are *n* (≥1) mice in both drug and vehicle groups, and (2) a 21-day trial with tumor volume measured at treatment initiation and then twice every week to produce 8 data points for every mouse. Drug efficacy is measured by how much drug treatment slows down tumor growth (β_2_/β_1_ in Equation 4). Power curves were obtained by computational simulations based on parameters obtained from fitting the cisplatin dataset by Equation 4 (Fig. 3c).

We observed that if the number of PDXs is the same, more mice per PDX confer better statistical power. For example, to achieve 80% power, we need about 28 PDXs for the 1:1 design, and 11 PDXs for the 3:3 design. More importantly, statistical power is comparable for designs with similar number of total mice. For example, when the drug efficacy is 20%, that is, the drug reduces tumor growth rate by 20%, the following designs all achieve 90% power at 0.05 significance level: 36 PDX with 1:1 design, 19 PDXs with 2:2 design, 13 PDXs with 3:3 design, 10 PDXs with 4:4 design, and so on. Therefore, in practice, we can use the 1:1 design if there are many PDXs at disposal—the 1×1×1 approach (Gao et al., 2015). Alternatively, we can increase the number of mice per PDX to boost power, especially when there is only a limited number of suitable PDXs, e.g., PDXs carrying a particular mutation or PDXs of a specific subtype.

We also observed that fewer PDXs are needed for a more potent drug to reach same statistical power. For example, to achieve 80% statistical power at 0.05 significance level by the 3:3 design, we need about 40, 11, and 5 PDXs for drugs with 10%, 20%, and 30% efficacy, respectively. When a drug is potent enough, all n:n designs achieve high power with very small number of PDXs. In such cases, we use a good number of PDXs not for statistical power but for better representation of tumor heterogeneity.

### Survival analysis in MCTs

In clinical trials, patient survival is usually assumed to be independent of each other. In MCTs, this assumption no longer holds because mice are now clustered within PDXs, and mice of same PDX tend to have more similar survival time, while their survival time between treatments is highly correlated (Fig. 4a). Further, PDXs can vary greatly in growth rate (or hazard) and drug response (**Fig. S9**). Therefore, we use an additive frailty model to model the heterogeneity on hazard and drug efficacy under the clustered population structure of MCTs (see Equation 5 in Materials and Methods). The additive frailty model is an extension of the Cox proportional hazards model wildly used in clinical trials. It has two frailty terms, the first one *u_i_* quantifies PDX growth rate heterogeneity and the second one *v_i_* measures drug response heterogeneity.

**Figure 4.**
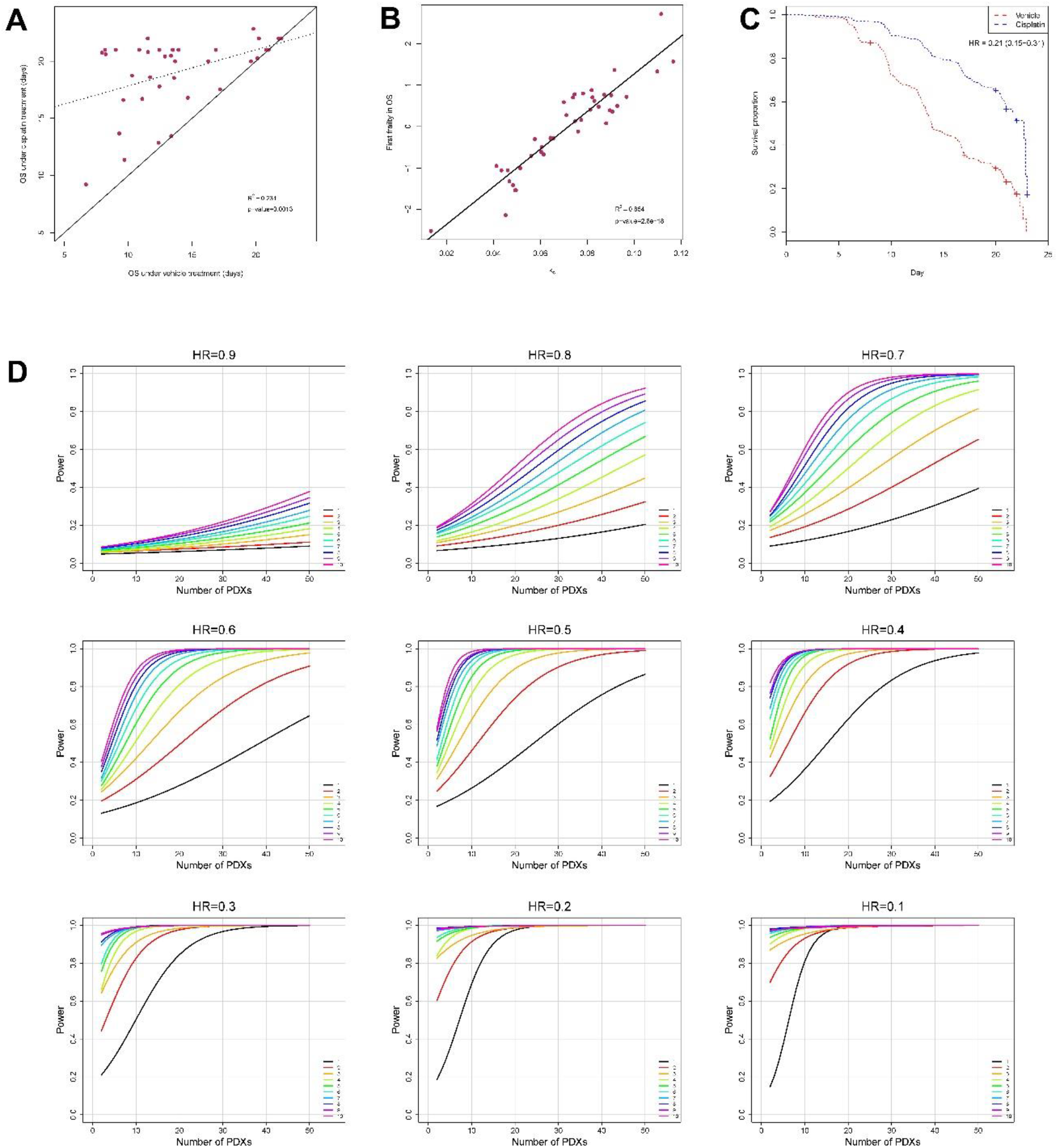
Survival analysis in a cisplatin MCT. (**a**) The median progression free survival (PFS) times of PDXs under cisplatin and vehicle treatment are highly correlated. The dotted line is the linear regression lines, and the solid line is a line with unit slope. (**b**) The first frailty term *u_i_* in Equation 5 is positively correlated with the tumor growth rate *k_c_*. (**c**) Survival curves under cisplatin and vehicle treatments. Additive frailty model gives more accurate hazard ratio (HR) than the Cox proportional hazards model whose estimation is 0.36 (95% CI: 0.28-0.46). (d) Statistical power curves at significance level α=0.05 when the hazard ratio is 0.9 to 0.1 for the survival analysis. The 10 colored curves in each graph denote the number of mice per PDX per arm.

We use the cisplatin MCT to illustrate the utilization of the additive frailty model. Overall survival (OS) is defined as tumor volume tripling time. We fit the cisplatin MCT dataset by Equation 5, and observed that both frailty terms are significant larger than 0 (Wald test p-value<0.05), proving that the PDXs grow at different rate and had different responses to cisplatin. In fact, the first frailty term *u_i_* is negatively correlated with tumor growth rate in the vehicle group, as expected (R^2^ = 0.85, Fig. 4b).

Drug efficacy can be estimated more accurately by excluding the influence of tumor growth heterogeneity and considering drug response heterogeneity, which is measured by the second frailty term *v_i_*. Indeed, the hazard ratio (HR) is estimated to be 0.21 (95% CI: 0.15-0.31), much smaller than that obtained from the Cox proportional hazards model, which gives HR=0.36 (95% CI: 0.28-0.46) (Fig. 4c). These results show that without considering PDX heterogeneity, drug effect can be severely misestimated.

We performed statistical power analysis for the survival analysis by assuming the n:n designs and using parameters estimated from the cisplatin MCT with Weibull hazard functions (Figure 4d). Like in LMMs, statistical power is similar for designs with similar total number of mice.

### Biomarker discovery in MCTs

Genomic correlation to cetuximab efficacy in solid tumors has been well documented (Bertotti et al., 2015), and we previously reported a MCT for a cohort of 20 gastric cancer PDXs and found that EGFR expression to be a predictive biomarker for cetuximab on gastric cancer (L. Zhang et al., 2013). The cohort is now expanded to 27 PDXs (**Fig. S10**). We observed a strong correlation between EGFR expression and drug efficacy measured by tumor growth inhibition or TGI (Fig. 5a). But EGFR is ranked 157^th^ out of all genes.

**Figure 5.**
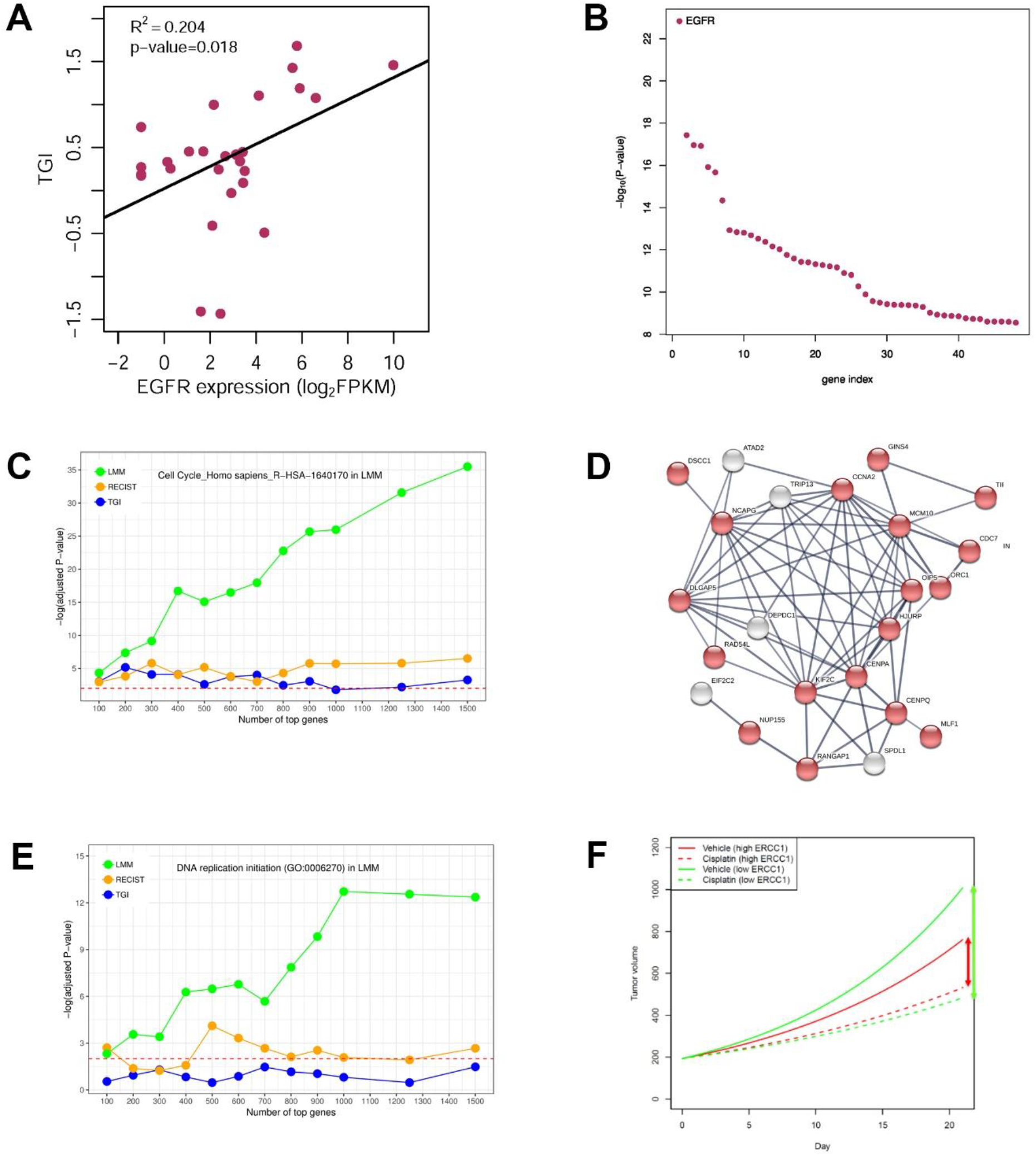
Biomarker discovery and MoA study in MCTs. (**a-b**): A MCT of 27 gastric cancer PDXs treated with cetuximab. EGFR is ranked 157^th^ among all genes based on Spearman rank correlation between EGFR expression and TGI (**a**), but is the top gene in predicting cetuximab efficacy based on a linear mixed model (LMM) (**b**). (**c-e**): A MCT of 16 PDXs treated with intraperitoneal injection of Irinotecan (100mg/kg, once per week for 2 to 3 weeks). (**c**): R-HAS-160170, the cell cycle pathway in Reactome2016 database, is consistently ranked as the most enriched pathway with 100 to 2000 top genes selected by a LMM, superior to top genes selected by methods based on categorical endpoints (e.g., RECIST, Table S2) and continuous endpoints (e.g., TGI). (**d**): DNA replication initiation (GO: 0006270) is the most enriched GO term based on top genes selected by the LMM. (**e**): A highly enriched protein-protein interaction network (p-value<10^-16^) consisted of 23 genes in the top 100 genes selected by the LMM. Red-colored nodes are ones involved in cell cycle (GO: 0007049). Dashed horizontal lines in (**c-d**) denotes p-value=0.01. (**f**): Mean tumor growth curves for PDXs with highest and lowest ERCC1 mRNA expression in a MCT of 21 gastric PDXs treated by cisplatin.

We used a LMM that explicitly models a gene’s effect on tumor growth to fit the efficacy data (Equation 6 in Materials and Methods). EGFR stands out as the most significant gene and its p-value, being1.5×10^-23^, is at least five orders of magnitude smaller than all other genes (Fig. 5b). EGFR as a predictive biomarker for cetuximab on gastric cancer is supported by a phase 2 clinical trial (X. Zhang et al., 2014) and a phase 3 clinical trial with data re-interpretation (**Fig. S11**) (Lordick et al., 2013). This study shows that simple analysis can produce many false positive hits to hamper biomarker discovery, especially when a drug target is unknown or there are off-target effects, while the more sophisticated LMM method can be superior in biomarker discovery.

### Mechanism of action study in MCTs

Investigating the relationship between efficacy and genomic data facilitates the study of a drug’s mechanism of action (MoA). Like in biomarker discovery, simple categorical and continuous endpoints, as a gross summery of efficacy, have various drawbacks. For example, the 4 categorical methods only measure efficacy in drug treatment group, ignoring the relative drug-to-vehicle efficacy. RTV ratio and TGI are dependent on calculation day and tumor growth rate (**Fig. S12**).

Irinotecan is a DNA topoisomerase I inhibitor that interrupts cell cycle in the S-phase by irreversibly arresting the replication fork, therefore causing cell death (Xu & Villalona-Calero, 2002). We conducted a MCT for 16 PDXs (**Fig. S13**), each PDX with 3 to 10 mice. We modeled the effect of gene expression on drug efficacy by a LMM (Equation 6). Top ranked genes were highly enriched for the cell cycle pathway R-HSA-160170 in the Reactome 2016 database (Fig. 5c), and for DNA replication initiation (Gene Ontology annotation GO: 0006270) (Fig. 5d), which perfectly reveals the MoA for irinotecan. A highly connected protein-protein interaction network for cell cycle is also identified from the 100 top ranked genes (Fig. 5e). In contrast, endpoint based methods are far less insightful (**Table S2-S4**, Fig. 5c-d).

### MCTs can explain paradoxical clinical trial results

Conflicting clinical trial reports exist regarding the role of ERCC1 expression in predicting cisplatin treatment on gastric cancer: some claimed that patients benefit more from low ERCC1 expression (De Dosso et al., 2013; Hirakawa et al., 2013; Kwon et al., 2007; Metzger et al., 1998; Miura et al., 2015), some stated the opposite (Baek et al., 2006; Bamias et al., 2010; Kim et al., 2011), while still others found no connection at all (Sonnenblick et al., 2012).

In a previous section, we described a cisplatin MCT which included 21 gastric cancer PDXs. We fit the tumor volume data by Equation 6. Parameter *β*_2_ quantifies how ERCC1 expression affects tumor growth when there is no drug intervention, as seen from the vehicle growth curves (Fig. 5f). Parameter *β*_4_ evaluates how ERCC1 expression impacts cisplatin’s efficacy on tumor growth, as seen by comparing the cisplatin growth curves with corresponding vehicle growth curves. These two parameters are at comparable magnitude but with opposite signs (*β*_2_ = −0.0155 and *β*_4_ = 0.0136). Therefore, when ERCC1 expression gets higher, tumor grows slower, but the benefit of cisplatin treatment is smaller as well (Fig. 5f).

In a clinical trial, patients with low/negative ERCC1 expression would have worse prognosis if they were not treated, and they could benefit more from cisplatin treatment. With treatment, their prognosis is improved, but whether it is better than the prognosis of ERCC1 high/positive patients is undetermined and depends on the trial population, hence we saw conflicting study conclusions.

## Discussion

MCTs are population-based efficacy trials mimicking human trials. Multiple mice are usually used per mouse model per arm to improve accuracy of efficacy measurement. For example, Bertotti et al. used 6 mice per PDX per arm in a two-arm MCT with 85 colorectal cancer PDXs to identify HER2 as a therapeutic target in Cetuximab-resistant colorectal cancers (Bertotti et al., 2011). It may also be feasible to use one mouse per model per arm when there is a large number of mouse models, which compensate the loss of measurement accuracy on individual mice (Gao et al., 2015; Murphy et al., 2016; Townsend et al., 2016; Williams, 2018). Caution must be exercised to use this approach though, when the number of mouse models is small, or high measurement accuracy of individual mouse models is mandated, or response varies greatly among mice of same mouse models, as commonly observed for immunotherapeutic agents on syngeneic models.

Our study established theoretic foundations for the design and analysis of MCTs. We first investigated tumor growth kinetics. Many complex mathematical models were used to describe tumor growth (Benzekry et al., 2014), but might not be particularly advantageous at the expense of more parameters and the need of more data points for model fitting. The exponential growth model is simple, interpretable and linear after a logarithmic transformation, and was shown to be adequate in most cases. Consequently, LMMs can describe nearly all MCTs, using quadratic terms of time if necessary.

We introduced additive frailty models to perform survival analysis for MCTs. The definition of PFS/OS can vary. For example, OS can be defined same as in human trials for leukemia PDXs (Townsend et al., 2016). For both LMMs and frailty models, we performed power simulations that give concrete recommendations on trial design. In particular, we answered the frequently asked questions on how many mouse models and how many mice per model to use, with flexible combination of the two numbers. MCTs can be asymmetric, i.e., unequal numbers of mice in arms. LMMs and frailty models are flexible for covariates, for example, a fixed effect for site can be incorporated if a MCT is conducted at multiple sites.

In conclusion, methods proposed in this study make the design and analysis of MCTs more rational and powerful.

## Materials and Methods

### Mouse models, studies and transcriptomic profiling

The establishment of mouse models and the conduct of mouse efficacy studies were described previously (Yang et al., 2013; Yang et al., 2016; L. Zhang et al., 2013). Briefly, for PDX models, freshly resected patient tumors were sliced into roughly 3×3×3mm^3^ chunks and engrafted subcutaneously on the flanks of immunocompromised mice (BALC/c, NOD/SCID, NOG, etc.). Tumor growth was monitored by a caliper twice a week to establish the first passage of a PDX model. Tumor was harvested for next round of engraftment when it reached 500-700 mm^3^ (1/2length×width^2^). A series of engraftment produced subsequent passages of the model. For CDX and syngeneic models, cell suspension (0.1-5×10^6^ cell/mouse) was injected into immunocompromised mice and immunocompetent mice (C57BC/6, BALB/c, etc.), respectively, to induce tumor. Pharmacological dosing started when a tumor was normally 100-300mm^3^, tumor volume was measured twice a week until the tumor was reaching 3000mm^3^, by then the mouse was euthanized. All animal studies were conducted at Crown Bioscience SPF facility under sterile conditions and were in strict accordance with the Guide for the Care and Use of Laboratory Animals of the National Institutes of Health. Protocols of all studies were approved by the Committee on the Ethics of Animal Experiments of Crown Bioscience, Inc. (Crown Bioscience IACUC Committee). Mouse models and cell lines were profiled by RNA-seq on Illumina HiSeq series platforms by certified service providers, as previously described (Guo et al., 2016).

### Categorical efficacy endpoints in mouse studies

Four categorical endpoint methods were evaluated, including the Response Evaluation Criteria In Solid Tumors (RECIST) criteria (Eisenhauer et al., 2009), a 3-category or 3-cat method (Bertotti et al., 2015), the 4-response mRECIST criterion (Gao et al., 2015), and a 5-category or 5-cat method (Houghton et al., 2007). Briefly, the RECIST-based criterion categorizes drug responses into complete response (CR), partial response (PR), stable disease (SD) and progressive disease (PD) based on relative tumor volume, or RTV, at a later day relative to treatment initiation day (CR: RTV=0, PR: 0<RTV≤0.657, SD: 0.657<RTV≤1.728, PD: RTV>1.728). Metastasis is not considered because it rarely occurs in subcutaneous implantation. The 3-cat method classifies response into PD, SD and objective response (OR) based RTV as well (OR: RTV≤0.65, PD: RTV≥1.35, SD: 0.65<RTV<1.35). The mRECIST method considers tumor growth kinetics 10 days after treatment initiation and classifies responses into CR, PR, SD and PD using two RTV-based quantities: best response and best average response. The 5-cat method classifies responses into maintained CR (MCR), CR, PR, SD and PD based on RTV (PD: RTV>0.50 during the study period and RTV>1.25 at end of study, SD: RTV>0.50 during the study period and RTV≤1.25 at end of study, PR: 0<RTV≤0.50 for at least one time point, CR: RTV=0 for at least one time point, MCR: RTV=0 at end of study). In the definitions of MCR and CR, we also use RTV=0 to designate disappearance of measurable tumor mass to replace the convention (TV<0.10cm^3^) used in Houghton et al., 2007. For all 4 methods, the admissive initial tumor volume is 50~300mm^3^.Objective response is defined as OR, CR+PR, MCR+CR+PR in the 3-cat, RECIST/mRECIST and 5-cat methods, respectively.

### Continuous efficacy endpoints in mouse studies

We briefly describe 4 continuous endpoints here. (a) Progression-free survival (PFS) is defined as tumor volume doubling time and obtained by linear intrapolation on tumor growth data. Specifically, if the PFS is between day *d*_1_ and day *d*_2_, then it is *d*_1_ + (*d*_2_ − *d*_1_)(2*TV*_0_ − *TV*_1_)/(*TV*_2_ − *TV*_1_) where *TV*_1_, *TV*_2_ and *TV*_0_ are tumor volumes at *d*_1_, *d*_2_ and treatment initiation day. (b) RTV ratio is the ratio of RTV between drug group and vehicle group at a specific day *d* and equals RTV_t_/RTV_c_, where RTV_t_ is the relative tumor volume between day *d* and treatment initiation day for the drug treatment group, and RTV_c_ is accordingly defined for the vehicle group. (c) Tumor growth inhibition (TGI) has several definitions, it can be defined as 1- RTV_t_/RTV_c_, or as 1-ΔT/ΔC where ΔT and ΔC are tumor volume changes relative to initial volume for drug group and vehicle group, respectively, at a specific day. (d) The ratio of growth rates between drug group and vehicle group is defined as *k*_t_/*k*_c_ where *k*_t_ and *k*_c_ are the growth rates obtained by modeling tumor growth data for the two groups by Equation 1. More general, we can introduce a new endpoint called AUC ratio, which reduces to ratio of growth rates when tumor grows under exponential kinetics (Fig. S5). Unique treatment models with at least 10 mice were used to calculate continuous endpoints, including 621 unique treated PDXs, 739 CDXs and 438 syngeneic models.

### Modeling tumor growth

Tumor growth under exponential kinetics is modeled by

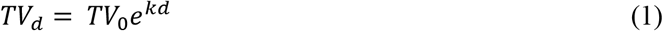

Where *TV_0_* is the initial tumor volume, *TV*_d_ is the tumor volume at day *d*, and *k* is the tumor growth rate. A logarithmic transformation gives

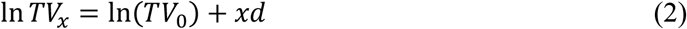

### Linear mixed models for the cisplatin dataset

A general model can be specified for tumor volume, in log scale, at day *t* for mouse *i* within PDX *j* as follows:

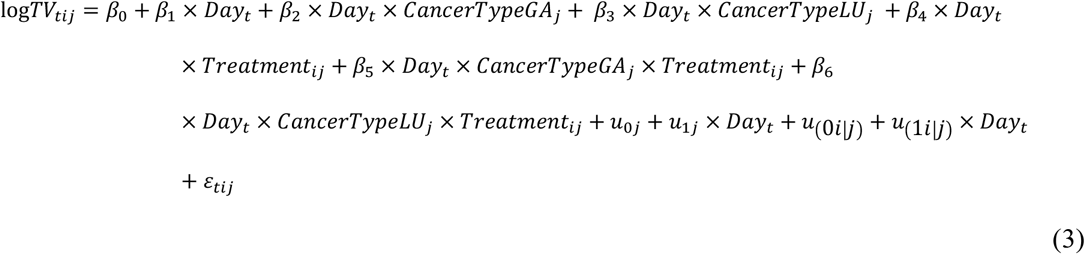

LU is lung cancer, GA is gastric cancer and ES is esophageal cancer. The model uses vehicle in ES as the reference. There are 6 fixed effects: *β*_0_ for the intercept, *β*_1_ for the time slope, *β*_2_ and *β*_3_ quantify the growth rate difference of GA and LU with respect to ES, *β*_4_ measures cisplatin effect, *β*_5_ and *β*_6_ measures if GA and LU respond differently to cisplatin. The model also has 5 random effects, including the residual *ε_tij_*. In a MCT, we view the cohort of PDXs as random samples from a PDX or patient population, therefore, they have different growth rates, which is modeled by random effect *u*_1j_ associated with the time slope. Similarly, we model growth difference for mice within a PDX by the random effect *u*_1i|j_. Mice and PDX may have different starting tumor volumes, modeled by the two random effects on intercept *u*_0j_ and *u*_0i|j_.

### Power calculation based on computational simulation

Power calculation was based on parameters (e.g., variance and covariance of random effects) estimated from fitting the cisplatin dataset by a LMM:

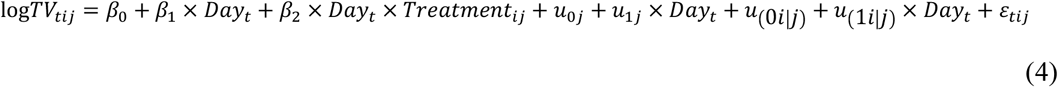

At significance level α=0.05, we obtained power curves by simulations for β_2_/β_1_= −0.1 to −0.9, that is, drug treatment reduces tumor growth rate by 10% to 90%.

### Additive frailty models for survival analysis

In the additive frailty model, the hazard function for the *j-*th mouse of the *i-*th mouse model is given by

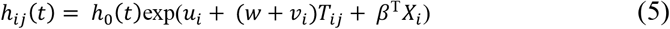

where *h*_0_(*t*) is the baseline hazard function. Parameter *u_i_* is the random effect (the first frailty term) associated with the *i-*th mouse model that captures its characteristic growth, thus survival behavior, without drug treatment. Parameter *v_i_* is the random effect (the second frailty term) associated with the *i*-th mouse model that depicts its drug response. Parameter *w* measures the drug treatment effect on all mouse models. *T*_*ij*_ is the treatment variable and equals 0 for the vehicle treatment and 1 for the drug treatment; *X*_*i*_ is a vector for the mouse model’s covariates, e.g., cancer type and genomic features; *β*^T^ is the parameter vector quantifying the fixed effects of the covariates. The two random effects *u_i_* and *v_i_* assume a bivariate normal distribution with zero means, variance *σ*^2^ and *τ*^2^, and covariance ρ*στ*. If the two random effects *u_i_* and *v_i_* are removed, the model reduces to the Cox proportional hazards model. Model fitting was done by the R package frailtypack (version 2.12.6), assuming Weibull distribution for the hazard function (Rondeau & Gonzalez, 2005).

### Linear mixed models for the biomarker discovery

The following LMM is used for single-gene biomarker discovery by fitting efficacy data from a MCT:

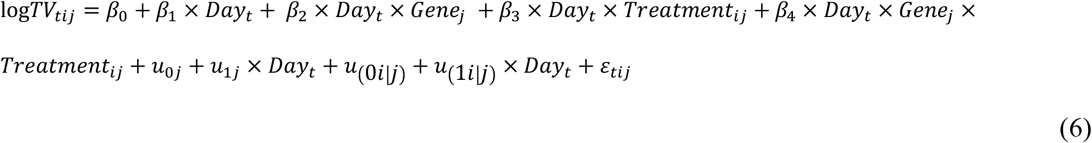

In this model, *Gene* is a covariate for the genomic status (expression, mutation, copy number variation, etc.) of a gene.

### Gene list enrichment analysis

A list of top ranked genes were used as input to the Enrichr web server (http://amp.pharm.mssm.edu/Enrichr/) for their enrichment in the “Reactome 2016” pathway database and in the “GO Biological Process 2018” database (Chen et al., 2013). Adjusted p-values were used to rank enriched pathways and biological processes.

### Protein-protein interaction network analysis

A list of top ranked genes were analyzed for protein-protein interactions in the STRING database (version 10.5 at https://string-db.org) (Szklarczyk et al., 2015). Default settings were used except the value for “minimum required interaction score” changed from “medium confidence (0.400)” to “high confidence (0.700)”.

## Acknowledgements

The authors would like to express their gratitude to the in vivo team members at the Translational Oncology Division of Crown Bioscience, Inc. for contributing all the in vivo efficacy data.

## Abbreviations

PDX: patient-derived xenograft
CDX: cell line-derived xenograft
GEMM: genetically engineered mouse model
MCT: mouse clinical trial
LMM: linear mixed model
RCT: randomized controlled trial
PFS: progression-free survival
OS: overall survival
ES: esophageal cancer
GA: gastric cancer
LU: lung cancer
TGI: tumor growth inhibition
EGFR: epidermal growth factor receptor
ERCC1: ERCC excision repair 1
HER2: human epidermal growth factor receptor 2
RECIST: Response Evaluation Criteria in Solid Tumors
HR: hazard ratio
OR: objective response
CR: complete response
PR: partial response
SD: stable disease
PD: progressive disease
RTV: relative tumor volume
MoA: mechanism of action.

